# SARS-CoV-2 Productively Infects Human Gut Enterocytes

**DOI:** 10.1101/2020.04.25.060350

**Authors:** Mart M. Lamers, Joep Beumer, Jelte van der Vaart, Kèvin Knoops, Jens Puschhof, Tim I. Breugem, Raimond B.G. Ravelli, J. Paul van Schayck, Anna Z. Mykytyn, Hans Q. Duimel, Elly van Donselaar, Samra Riesebosch, Helma J.H. Kuijpers, Debby Schipper, Willine J. van de Wetering, Miranda de Graaf, Marion Koopmans, Edwin Cuppen, Peter J. Peters, Bart L. Haagmans, Hans Clevers

**Author notes:** Equal contribution (Mart M. Lamers, Joep Beumer, Jelte van der Vaart). Equal contribution (Bart L. Haagmans, Hans Clevers).

## Abstract

COVID-19, caused by SARS-CoV-2, is an influenza-like disease with a respiratory route of transmission, yet clinical evidence suggests that the intestine may present another viral target organ. Indeed, the SARS-CoV-2 receptor angiotensin converting enzyme 2 (ACE2) is highly expressed on differentiated enterocytes. In human small intestinal organoids, enterocytes were readily infected by SARS-CoV and SARS-CoV-2 as demonstrated by confocal- and electron-microscopy. Consequently, significant titers of infectious viral particles were measured. mRNA expression analysis revealed strong induction of a generic viral response program. We conclude that intestinal epithelium supports SARS-CoV-2 replication.

**One Sentence Summary:** SARS-CoV-2 infection of enterocytes in human small intestinal organoids

## Main Text

Severe Acute Respiratory Syndrome (SARS), caused by the coronavirus SARS-CoV, emerged in 2003 (*1*). Late last year, a novel transmissible coronavirus (SARS-CoV-2) was noted to cause an influenza-like disease ranging from mild respiratory symptoms to severe lung injury, multi-organ failure and death (*2–4*). SARS-CoV-2 belongs to the *Sarbecovirus* subgenus (genus *Betacoronavirus,* family *Coronaviridae),* together with SARS-CoV (*5–7*). Angiotensin-converting enzyme 2 (ACE2) is the SARS-CoV receptor (*8, 9*). The spike proteins of the two viruses both bind to ACE2, while soluble ACE2 blocks infection by SARS-CoV-2 (*10–13*). Transmission of SARS-CoV-2 is thought to occur through respiratory droplets and fomites. In accordance, the virus can be detected in upper respiratory tract samples, implicating the nasopharynx as site of replication. In human lung, ACE2 is expressed mainly in alveolar epithelial type II cells and ciliated cells (*14–16*). Yet, highest expression of ACE2 in the human body occurs in the brush border of intestinal enterocytes (*14, 17*). Even though respiratory symptoms dominate the clinical presentation of COVID-19, gastrointestinal symptoms are observed in a subset of patients (*18,19*). Moreover, viral RNA can be found in rectal swabs, even after nasopharyngeal testing has turned negative, implying gastro-intestinal infection and a fecal–oral transmission route (*20–22*). Here, we have utilized human small intestinal organoids (hSIOs) to study infection by SARS-CoV-2.

We first demonstrated that SARS-CoV and SARS-CoV-2 readily infect organoid-derived human airway epithelium cultured in 2D (Fig 1A). The viruses targeted ciliated cells, but not goblet cells (Fig. 1B-C). hSIOs are established from primary gut epithelial stem cells, can be expanded indefinitely in 3D culture and contain all proliferative and differentiated cell types of the *in vivo* epithelium (*23*). Of note, hSIOs have allowed the first *in vitro* culturing of Norovirus (*24*). We exposed ileal hSIOs grown under four different culture conditions to SARS-CoV and SARS-CoV-2 at a multiplicity of infection (MOI) of 1. hSIOs grown in Wnt-high expansion medium (EXP) overwhelmingly consist of stem cells and enterocyte progenitors. Organoids grown in differentiation medium (DIF) contain enterocytes, goblet cells and low numbers of enteroendocrine cells (EECs). Addition of BMP2/4 to DIF (DIF-BMP) leads to further maturation (*25*). In the final condition, we induced expression of NeuroG3 with doxycycline to raise EECs numbers (JB and HC, unpublished; Suppl Fig 3D).

**Figure 1.**
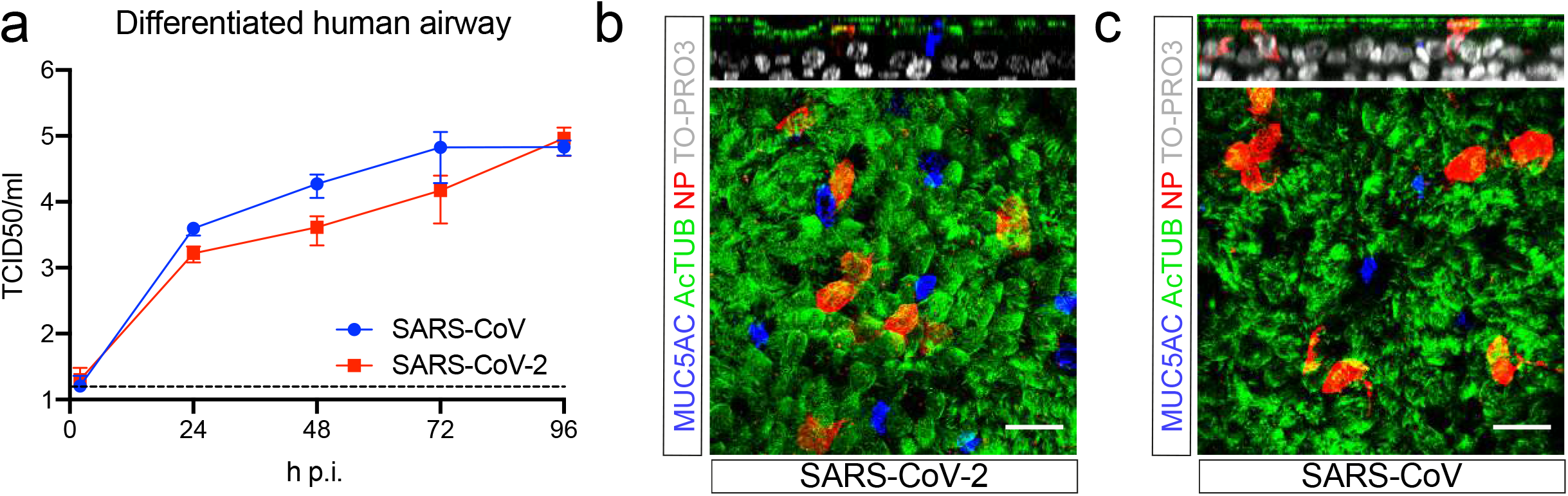
SARS-CoV and SARS-CoV-2 infect 2D human airway cultures. a) Live virus titers can be observed by virus titrations on VeroE6 cells of apical washes at 2, 24, 48, 72 and 96h after infection with SARS-CoV (blue) and SARS-CoV-2 (red). The dotted line indicates the lower limit of detection. Error bars represent SEM. N=4. b and c) Immunofluorescent staining of SARS-CoV-2 (b) and SARS-CoV (c) infected differentiated airway cultures. Nucleoprotein (NP) stains viral nucleocapsid (red), which colocalized with the ciliated cell marker AcTUB (green). Goblet cells are identified by MUC5AC (blue). Nuclei are stained with TO-PRO3 (white). Scale bars indicate 20μM.

Samples were harvested at multiple timepoints post infection and processed for the analyses given in Figs 2–5. Both SARS-CoV and SARS-CoV-2 productively infected hSIOs as assessed by qRT-PCR targeting the E gene and by live virus titrations on VeroE6 cells (Fig. 2 for lysed organoids; Fig. S1 for organoid supernatant). Infectious virus particles and viral RNA increased to significant titers for both viruses in all conditions. As EXP medium supported virus replication (Fig. 2A, E), enterocyte progenitors appeared to be a primary viral target. Differentiated organoids (DIF; DIF-BMP) produced slightly (non-statistically significant) lower levels of infectious virus (Fig. 2; Fig. S1). In organoids induced to generate EECs, virus yields were similar to those in EXP medium (Fig. 2D, H). In differentiated hSIOs, SARS-CoV-2 titers remained stable at 60 hours post infection, while SARS-CoV titers dropped 1-2 log (Fig. 2B, C, F, G). The latter decline was not observed in infected hSIOs grown in EXP. Culture supernatants across culture conditions contained lower levels of infectious virus compared to lysed hSIOs, implying that virus was primarily secreted apically (Fig. S1A-D). Despite this, viral RNA was detected readily in culture supernatants correlating with the infectious virus levels within hSIOs (Fig. 2E-H; Fig. S1 E-H).

**Figure 2.**
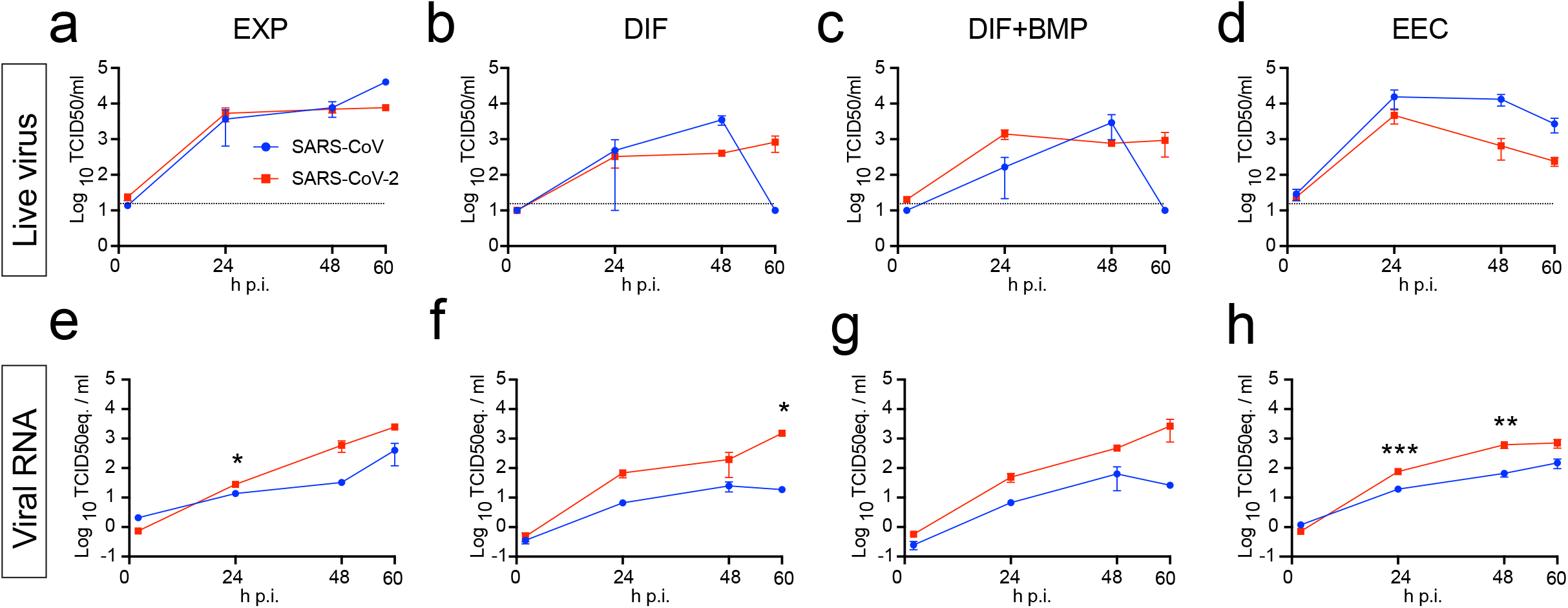
SARS-CoV and SARS-CoV-2 replicate in hSIOs. a-d) Live virus titers can be observed by virus titrations on VeroE6 cells of lysed organoids at 2, 24, 48 and 60h after infection with SARS-CoV (blue) and SARS-CoV-2 (red). Different medium compositions show similar results. e-h) qPCR analysis targeting the E gene of similar timepoints and medium compositions as a-d. The dotted line indicates the lower limit of detection. Error bars represent SEM. N=3.

ACE2 mRNA expression differed greatly between the four conditions. EXP-hSIOs express 300-fold less ACE2 mRNA compared to DIF-hSIOs, when analyzed in bulk (Fig. S2). BMP treatment induced 6.5-fold upregulation of ACE2 mRNA compared to DIF treatment alone. Since this did not yield infection rate differences, the DIF-BMP condition was not analyzed further.

To determine the target cell type, we then performed confocal analysis on hSIOs cultured in EXP, DIF, or EEC conditions. We stained for viral dsRNA, viral nucleocapsid protein, KI67 to visualize proliferative cells, actin (using phalloidin) to visualize enterocyte brush borders, DNA (DAPI) and cleaved caspase 3 to visualize apoptotic cells. Generally, comparable rates of viral infections were observed in the organoids growing under the three remaining conditions. We typically noted staining for viral components (white) in rare, single cells at 24 hours. At 60 hours, the number of infected cells had dramatically increased (Fig. 3A).

**Figure 3.**
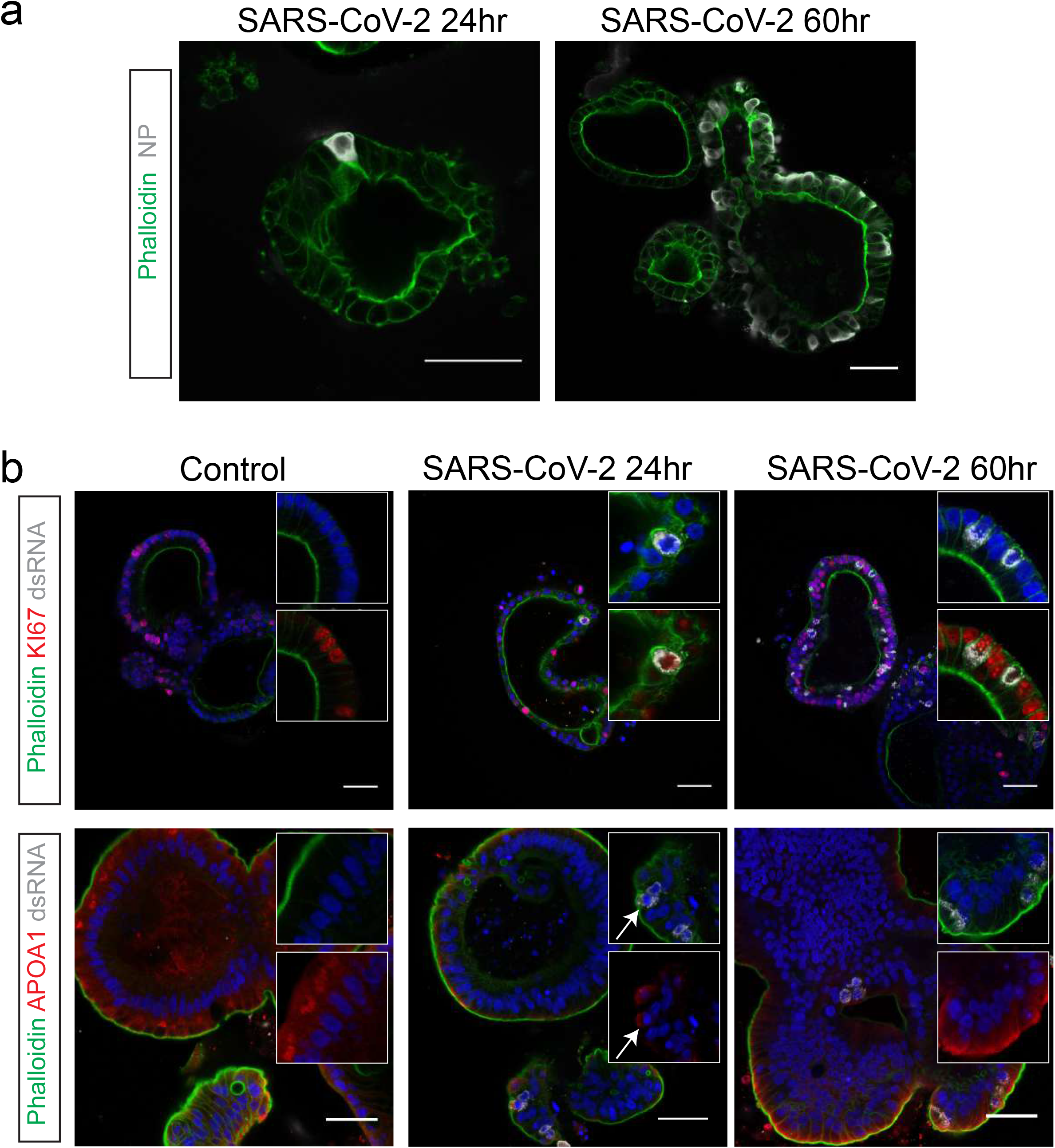
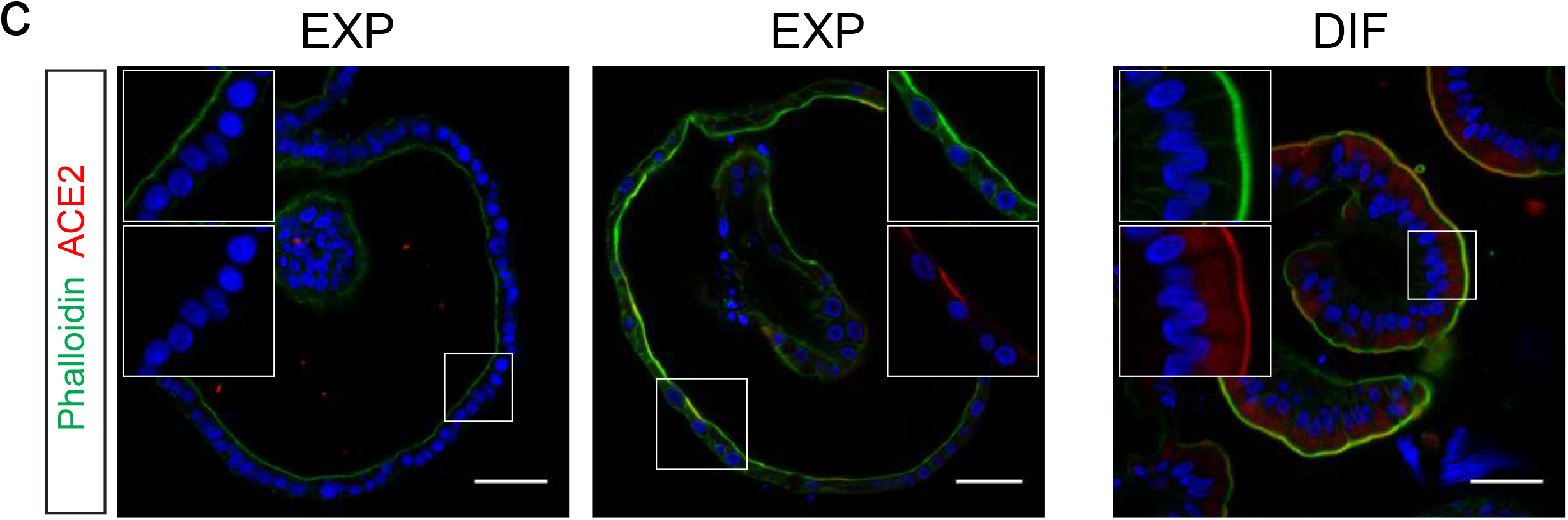
SARS-CoV-2 infects proliferating cells and enterocytes. a) Immunofluorescent staining of SARS-CoV-2-infected intestinal organoids. Nucleoprotein (NP) stains viral capsid. After 24 hours, single virus-infected cells are generally observed in organoids. These small infection clusters spread through the whole organoid after 60 hours. b) SARS-CoV-2 infects both post-mitotic enterocytes identified by Apolipoprotein A1 (APOA1) and dividing cells that are KI67-positive. Infected cells are visualized by dsRNA staining. Enterocytes are shown in differentiated organoids, and proliferating cells in expanding organoids. Arrows point to APOA1-positive cells. c) Immunofluorescent staining of ACE2 in intestinal organoids in expansion and differentiation condition. All scale bars are 50 μm.

Infected cells invariably displayed proliferative enterocyte progenitor-phenotypes (EXP, Fig. 3B, top) or ApoA1+ enterocyte-phenotypes (DIF, Fig. 3B, bottom). Of note, SARS-CoV also readily infected enterocyte precursors (Fig. S3A-B) as shown previously (*26, 27*). Some infected enterocyte progenitors were in mitosis (Fig. S3C). While EEC-organoids produced appreciable titers, we never observed infection of Chromogranin-A+ EECs (Fig. S3D-E). We also did not notice infection of Goblet cells across culture conditions. At 60 hours, apoptosis became prominent in both SARS-CoV and SARS-CoV-2 infected enterocytes (Fig. S5). ACE2 protein was readily revealed as a bright and ubiquitous brush border marker in hSIOs in DIF medium (Fig. 3C). In hSIOs in EXP medium, ACE2 staining was much lower -yet still apical-in occasional cells in a subset of organoids that displayed a more mature morphology (Fig. 3C). In immature (cystic) organoids within the same cultures, the ACE2 signal was below the detection threshold. Percentage of infected organoids under EXP and DIF conditions are given in Fig S4. Fig. S5 gives images and quantification of apoptotic cells upon infection.

Unsupervised transmission electron microscopy (TEM) (*26*) was performed on selected highly infected samples. Figure 4 shows two hSIOs, selected from 42 imaged hSIOs, 60 h post SARS-CoV-2 infection. These differ in the state of infection: whereas the cellular organization within organoid 1 was still intact (Fig. 4A entire organoid; intermediate magnification B-D; high magnification E-K), many disintegrated cells can be seen in organoid 2 (Fig. 4 Bottom (Fig. 4L entire organoid; intermediate magnification 4M-O; high magnification 4P-R). Viral particles of 80-120 nm occurred in the lumen of the organoid (4I), at the basolateral (4J) and apical side (4K) of enterocytes. The double-membrane vesicles which are the subcellular site of viral replication (*27*) are visualized in Figure 4E and 4P. The nuclei in both organoids differed from nuclei in mock-infected organoids by a slightly rounder shape. The nuclear contour index (*28*) was 4.0+/− 0.5 vs 4.3+/− 0.5 for control set. There was more heterochromatin (4N), and one or two dense nucleoli in the center (4O).

**Figure 4.**
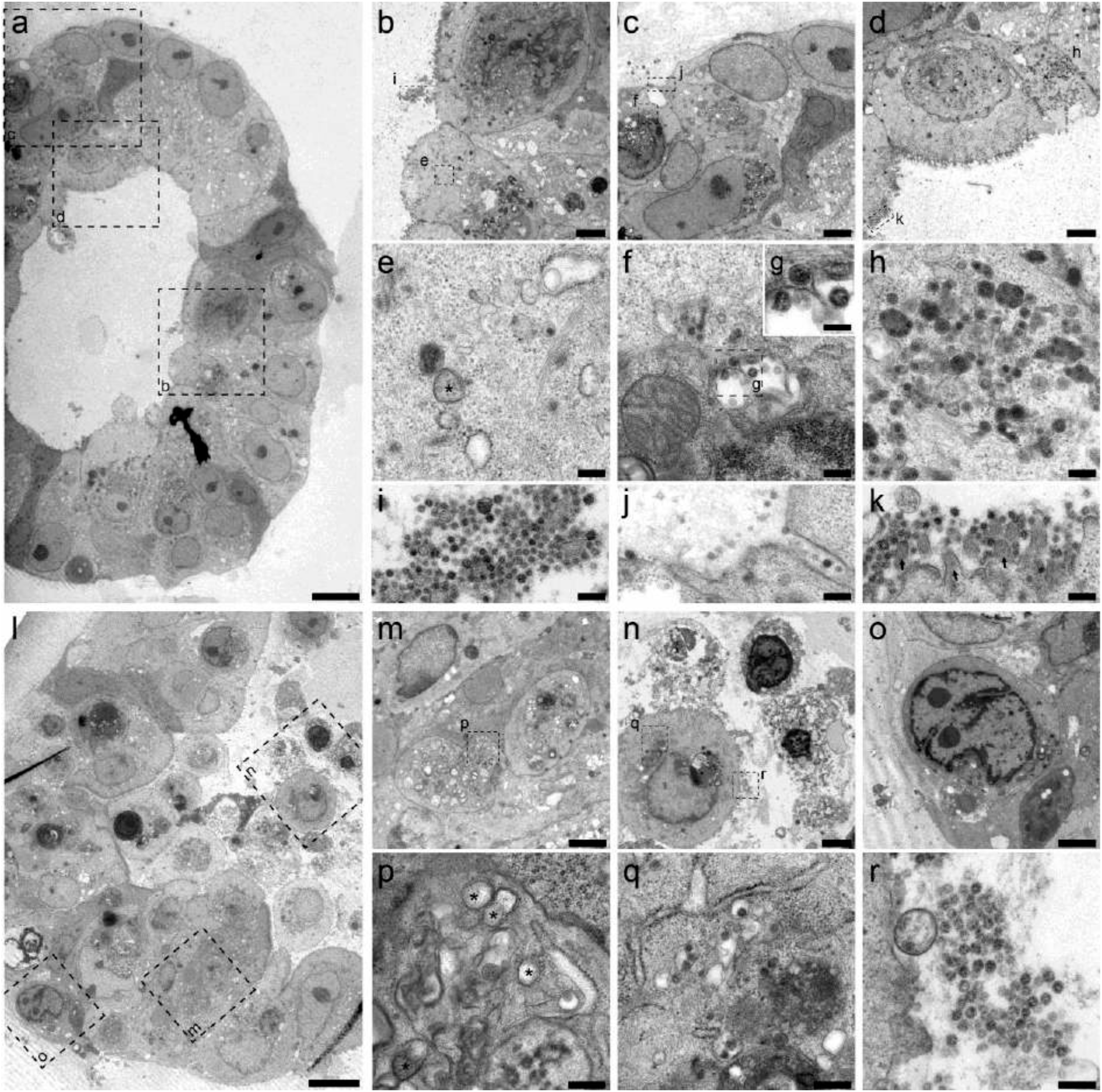
Transmission electron microscopy analysis of SARS-CoV-2 infected intestinal organoids. a) Overview of an intact organoid (a) showing the onset of virus infection (b-d) at different stages of the viral lifecycle, i.e. early double membrane vesicles (DMVs) (e; asterisk), initial viral production in the Golgi apparatus (f-g) and complete occupation of virus particles inside the endomembrane system (h). Extracellular viruses are observed in the lumen of the organoid (i), and are found at the basal (j) and apical side (k) alongside the microvilli (arrows). Scale bars represent 10 μm (a), 2.5 μm (b-d), 250 nm (e, f, h-k) & 100 nm (g). 1) Overview of an organoid (l) showing severely infected cells (m,o), disintegrated cells (o) and stressed cells as evident from the atypical nucleoli (p). Intact cells reveal DMV areas of viral replication (p; asterisks) and infected Golgi apparatus (q). Extracellular clusters of viruses are shown in (r). Scale bars represent 10 μm (l), 2.5 μm (m-p) & 250 nm (p-r). Data was deposited to the Image Data Resource (https://idr.openmicroscopy.org) under accession number idr0083.

We then performed mRNA sequence analysis to determine gene expression changes induced by SARS-CoV and SARS-CoV-2-infection of hSIOs cultured continuously in EXP medium and hSIOs cultured in DIF medium. Infection with SARS-CoV-2 elicited a broad signature of cytokines and interferon stimulated genes (ISGs) attributed to type I and III interferon responses (Fig 5A, Supplementary table 1-2), as confirmed by Gene Ontology analysis (Fig. 5B). An overlapping list of genes appeared in SARS-CoV-2-infected DIF organoids (Fig. S6, Supplementary table 3). RNA sequencing analysis confirmed differentiation of DIF organoids into multiple intestinal lineages, including ACE2 upregulation (Fig. S7). SARS-CoV also induced ISGs, yet to a much lower level (Supplementary table 4). Fig 5C visualizes the regulation of SARS-CoV-2-induced genes in SARS-CoV infected organoids. This induction was similar to infections with other viruses like norovirus (*29*), rotavirus (*30*) and enteroviruses (*31, 32*). A recent study (*33*) describes an antiviral signature induced in human cell lines after SARS-CoV-2 infection. While the ISG response is broader in intestinal organoids, the induced gene sets are in close agreement between the two datasets (Fig. S8). One striking similarity was the low expression of Type I and III interferons: we only noticed a small induction of the Type III interferon IFNL1 in SARS-CoV-2 infected organoids. In SARS-CoV-infected organoids, we did not observe any type I or type III interferon induction. We confirmed these findings by ELISA on culture supernatant and qRT-PCR, which in addition to IFNL1 picked up low levels of type I interferon IFNB1 in SARS-CoV-2 but not in SARS-CoV infected organoids (Fig. S9). The specific induction of IP-10/CXCL10 and ISG15 by SARS-CoV-2 was also confirmed by ELISA and qRT-PCR, respectively (Fig. S10). As in (*33*), a short list of cytokine genes was induced by both viruses albeit it to modest levels. For a comparison with (*33*), see Fig S11. Altogether these data indicate that SARS-CoV-2 induces a stronger interferon response than SARS-CoV in HIOs.

**Figure 5.**
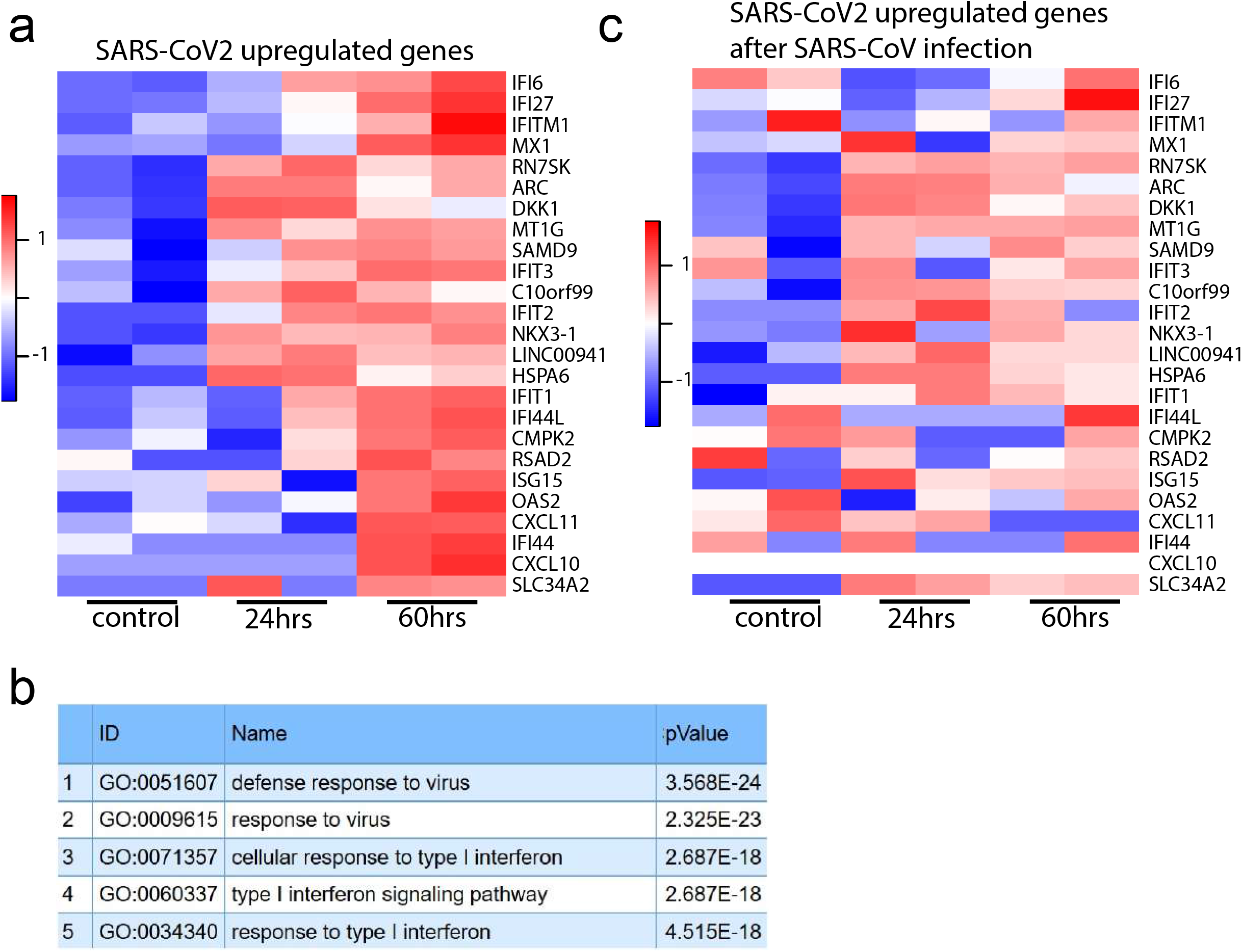
Transcriptomic analysis of SARS-CoV-2 infected intestinal organoids. a) Heatmaps depicting the 25 most significantly enriched genes upon SARS-CoV-2 infection in expanding intestinal organoids. Colored bar represents Z-score of log2 transformed values b) GO term enrichment analysis for biological processes of the 50 most significantly upregulated genes upon SARS-CoV-2 infected in intestinal organoids. c) Heatmaps depicting the genes from (a) in SARS-CoV infected expanding organoids. Colored bar represents Z-score of log2 transformed values.

Finally, the infection was repeated in a second experiment in the same ileal HIO line and analyzed after 72 hours. Analysis involved viral titration (Fig S12), confocal imaging (Fig S13) and RNA sequencing (Fig S14). This experiment essentially confirmed the observations presented above. A limited, qualitative experiment applying confocal analysis demonstrated infectability of two other lines available in the lab (one ileal, one duodenal) from independent donors (Fig S13).

This study shows that SARS-CoV and SARS-CoV-2 infect enterocyte lineage cells in a human intestinal organoid model. It appears of interest that we see at least equal infection rates of enterocyte-precursors and enterocytes whereas ACE2 expression increases ~1000-fold upon differentiation at the mRNA level (Fig. S2). This suggests that low levels of ACE2 may be sufficient for viral entry. The scale of the current SARS-CoV-2 pandemic suggests that this virus may continue to circulate in humans, causing yearly seasonal outbreaks. The fact that SARS-CoV-2 is the third highly pathogenic coronavirus (after SARS-CoV and MERS-CoV) to jump to humans within less than 20 years suggests that novel zoonotic coronavirus spillovers are likely to occur in the future. Despite this, limited information is available on coronavirus pathogenesis and transmission. This is in part due to the lack of *in vitro* cell models that accurately model host tissues. Very recently, it was already shown that human iPS cells differentiated towards a kidney fate support replication of SARS-CoV-2 (*13*). Our data imply that human organoids represent faithful experimental models to study the biology of coronaviruses.

## Acknowledgements

We thank Evelien Eenjes and Robbert Rottier for providing human lung material, Anko de Graaff and Hubrecht Imaging Centre (HIC) for microscopy assistance and Single Cell Discoveries for RNA library preparation, and the Utrecht Sequencing Facility (subsidized by the University Medical Center Utrecht, Hubrecht Institute, Utrecht University and NWO project 184.034.019).

## Author contributions

ML, JB and JV performed experiments and designed the study. KK & JQD prepared samples. KK & RBGR performed imaging. KK, JPvS, PJP & RGBR interpreted results. TB, AM, SR, DS and MG measured virus titers. JP analyzed RNAseq data. EC performed sequencing. MK, BH and HC supervised.

## List of Supplementary Materials

### Materials and Methods

#### Viruses and cell lines

Vero E6 cells were maintained in Dulbecco’s modified Eagle’s medium (DMEM, Gibco) supplemented with 10% fetal calf serum (FCS), HEPES, sodium bicabonate, penicillin (10,000 IU/mL) and streptomycin (10,000 IU/mL) at 37°C in a humidified CO2 incubator. SARS-CoV (isolate HKU 39849, accession no. AY278491) and SARS-CoV-2 (isolate BetaCoV/Munich/BavPat1/2020; European Virus Archive Global #026V-03883; kindly provided by Dr. C. Drosten) were propagated on Vero E6 (ATCC^®^ CRL 1586™) cells in Opti-MEM I (1X) + GlutaMAX (Gibco), supplemented with penicillin (10,000 IU/mL) and streptomycin (10,000 IU/mL) at 37°C in a humidified CO2 incubator. The SARS-CoV-2 isolate was obtained from a clinical case in Germany, diagnosed after returning from China. Stocks were produced by infecting Vero E6 cells at a multiplicity of infection (MOI) of 0.01 and incubating the cells for 72 hours. The culture supernatant was cleared by centrifugation and stored in aliquots at −80°C. Stock titers were determined by preparing 10-fold serial dilutions in Opti-MEM I (1X) + GlutaMAX. Aliquots of each dilution were added to monolayes of 2 × 104 Vero E6 cells in the same medium in a 96-well plate. Twenty-four replicates were performed per virus stock. Plates were incubated at 37°C for 5 days and then examined for cytopathic effect. The TCID50 was calculated according to the method of Spearman & Kärber. All work with infectious SARS-CoV and SARS-CoV-2 was performed in a Class II Biosafety Cabinet under BSL-3 conditions at Erasmus Medical Center.

#### Cell culture of human intestinal organoids

Tissues from the duodenum and ileum was obtained from the UMC Utrecht with informed consent of the patient. All patients were males that were operated for resection of an intestinal tumor. A sample from non-transformed, normal mucosa was taken for this study. The study was approved by the UMC Utrecht (Utrecht, The Netherlands) ethical committee and was in accordance with the Declaration of Helsinki and according to Dutch law. This study is compliant with all relevant ethical regulations regarding research involving human participants.

Human small intestinal cells were isolated, processed and cultured as described previously (*23, 25*). Instead of Wnt conditioned media, the medium was supplemented with Wnt surrogate (0,15 nM, U-Protein Express). This was termed expansion (EXP) medium. For differentiation towards EECs, organoids were treated with 1 μg/ml doxycycline (Sigma) in ‘ENR’ medium (Beumer et al., in press). General differentiation (termed DIF) was achieved in ‘ENR’. BMP activation was achieved by withdrawing Noggin from ‘ENR’ and addition of BMP-2 (Peprotech, 50 ng/ml) and BMP-4 (Peprotech, 50 ng/ml) (termed DIF+BMP).

#### Human airway organoid culture and differentiation

Adult human lung stem cells were derived from non-tumour lung tissue obtained from patients undergoing lung resection. Lung tissue was obtained from residual, tumor-free, material obtained at lung resection surgery for lung cancer. The Medical Ethical Committee of the Erasmus MC Rotterdam granted permission for this study (METC 2012-512). Isolation of human bronchial airway stem cells was performed using a protocol adapted from Sachs and colleagues (2019). Bronchial tissues of adult lungs were cut into ~4mm sections, washed in Advanced DMEM supplemented with GlutaMax (Thermo Fisher), HEPES, penicillin (10,000 IU/mL) and streptomycin (10,000 IU/mL) (AdDF+++), and incubated with dispase (Corning) mixed with airway organoid (AO) medium (*34*) at a 1:1 ratio for 1 hour at 37°C. The digested tissue suspension was sequentially sheared using 10- and 5-ml plastic and flamed glass Pasteur pipettes. After shearing and filtering, lung cell pellets were then resuspended in 200 μL of growth factor reduced Matrigel (Corning) and plated in ~30 μL droplets in a 48 well tissue culture plate. Plates were placed at 37 degrees Celsius with 5% CO2 for 20 min to solidify the Matrigel droplets upon which 250uL of media was added to each well. Plates were incubated under standard tissue culture c

To obtain differentiated organoid-derived cultures, organoids were dissociated into single cells using TrypLE express. Cells were seeded on Transwell membranes (Corning) coated with rat tail collagen type I (Fisher Scientific). Single cells were seeded in AO growth medium: complete base medium (CBM; Stemcell Pneumacult-ALI) at a 1:1 ratio. After 2-4 days, confluent monolayers were cultured at air-liquid interphase in CBM. Medium was changed every 5 days. After 4 weeks of differentiation, cultures were used for infection experiments.

#### SARS-CoV and SARS-CoV-2 Infection

Organoids were harvested in cold AdDF+++, washed once to remove BME, and sheared using a flamed Pasteur pipette in AdDF+++ or TrypLE express, for undifferentiated and differentiated organoids, respectively. After shearing, organoids were washed once in AdDF+++ before infection at a multiplicity of infection (MOI) of 1 in expansion or differentiation medium. After 2 hours of virus adsorption at 37°C 5% CO2, cultures were washed twice with excess AdDF+++ to remove unbound virus. Organoids were re-embedded into 30 μL BME in 24-well tissue culture plates and cultured in 500 μL expansion or differentiation medium at 37°C with 5% CO2. Each well contained ~100,000 cells per well. Samples were taken at indicated timepoints by harvesting the medium in the well (supernantant) and the cells by resuspending the BME droplet containing organoids into 500 μL AdDF+++. Samples were stored at −80°C, a process which lysed the organoids, releasing their contents into the medium. 2D differentiated airway cultures were washed twice with 200 ul AdDF+++ before inoculation from the apical side at a MOI of 0.1 in 200 μL AdDF+++ per well. Next, cultures were incubated at 37°C 5% CO2 for 2 hours before washing 3 times in 200 μL AdDF+++. At the indicated timepoints, virus was collected from the cells by adding 200 μL AdDF+++ apically, incubating 10 min at 37°C 5% CO2, and storing the supernatant at −80°C. Prior to determining the viral titer, all samples were centrifuged at 2,000 g for 5 min. Live virus titers were determined using the Spearman & Kärber TCID50 method on VeroE6 cells. All work was performed in a Class II Biosafety Cabinet under BSL-3 conditions at Erasmus Medical Center.

#### Determination of virus titers using qRT-PCR

Supernatant and organoid samples were thawed and centrifuged at 2,000 g for 5 min. Sixty μL supernatant was lysed in 90 μL MagnaPure LC Lysis buffer (Roche) at RT for 10 min. RNA was extracted by incubating samples with 50 μL Agencourt AMPure XP beads (Beckman Coulter) for 15 min at RT, washing beads twice with 70% ethanol on a DynaMag-96 magnet (Invitrogen) and eluting in 30 μL MagnaPure LC elution buffer (Roche). Viral titers (TCID50 equivalants per mL) were determined by qRT-PCR using primers targeting the E gene (*35*) and comparing the Ct values to a standard curve derived from a virus stock titrated on VeroE6 cells.

#### Fixed immunofluorescence microscopy on differentiated human airway

Transwell inserts were fixed in 4% paraformaldehyde for 20 min, permeabilized in 70% ethanol, and blocked for 60 min in 10% normal goat serum in PBS (blocking buffer). Cells were incubated with primary antibodies overnight at 4°C in blocking buffer, washed twice with PBS, incubated with corresponding secondary antibodies Alexa350-, 488- and 594-conjugated antirabbit and anti-mouse (1:400; Invitrogen) in blocking buffer for 2 h at room temperature, washed two times with PBS, incubated with indicated additional stains (TO-PRO3), washed twice with PBS, and mounted in Prolong Antifade (Invitrogen) mounting medium. Viral nucleocapsid was stained with anti-NP (1:200; Sino biological), ciliated cells were stained with anti-AcTUB (1:100; Santa Cruz, 6-11B-1) and goblet cells with anti-MUC5AC (1:100; Invitrogen, 45M1) Samples were imaged on a LSM700 confocal microscope using ZEN software (Zeiss).

#### Immunostaining of hSIOs

Organoids were stained as described before (*25*). Primary antibodies used were goat anti-chromogranin A (1:500; Santa Cruz), mouse anti-nucleoprotein (1:200; Sino Biological), mouse anti-dsRNA (1:200;Scicons), mouse anti-APOA1 (1:100; Thermofisher, PA5-88109), rabbit anti-cleaved caspase 3 (1:200; Cell Signaling Technology, 9661), goat anti-ACE2 (1:100; R&D Systems, AF933) and rabbit anti-KI67 (1:200; Abcam, ab16667). Organoids were incubated with the corresponding secondary antibodies Alexa488-, 568-and 647-conjugated anti-rabbit and anti-mouse (1: 1,000; Molecular Probes) in blocking buffer containing 4’,6-diamidino-2-phenylindole (DAPI; 1;1,000, Invitrogen) and Phalloidin-Alexa488 (Thermofisher). Sections were embedded in Vectashield (Vector Labs) and imaged using a Sp8 confocal microscope (Leica). Image analysis was performed using ImageJ software.

#### Quantitative PCR

Quantitative PCR (qPCR) analysis was performed using biological and technical duplicates as described before (*36*). Primers were designed using the NCBI primer design tool, tested using a standard curve, and include:

ACE2_fw CGAAGCCGAAGACCTGTTCTA
ACE2_rev GGGCAAGTGTGGACTGTTCC
GAPDH_fw GGAGCGAGATCCCTCCAAAAT
GAPDH_rev GGCTGTTGTCATACTTCTCATGG

For qPCR analysis of IFNB1, IFNL1 and ISG15 we used RNA extracted from lysed organoids as described above. We used the following ThermoFisher TaqMan primer-probe mixes:

IFNB1: Hs00277188-s1
IFNL1: Hs00601677-g1
ISG15: Hs01921425-s1

Beta actin was amplified using the following primer set:

Bact_fw GGCATCCACGAAACTACCTT, Bact_rev AGCACTGTGTTGGCGTACAG

Bact_probe FAM-ATCATGAAGTGTGACGTGGACATCCG-BHQ1.

#### Bulk RNA sequencing

Library preparation was performed at Single Cell Discoveries (Utrecht, The Netherlands), using an adapted version of the CEL-seq protocol. In brief:

Total RNA was extracted using the standard TRIzol (Invitrogen) protocol and used for library preparation and sequencing. mRNA was processed as described previously, following an adapted version of the single-cell mRNA seq protocol of CEL-Seq (*37, 38*). In brief, samples were barcoded with CEL-seq primers during a reverse transcription and pooled after second strand synthesis. The resulting cDNA was amplified with an overnight In vitro transcription reaction. From this amplified RNA, sequencing libraries were prepared with Illumina Truseq small RNA primers. Paired-end sequencing was performed on the Illumina Nextseq500 platform using barcoded 1 x 75 nt read setup. Read 1 was used to identify the Illumina library index and CEL-Seq sample barcode. Read 2 was aligned to the CRCh38 human RefSeq transcriptome, with the addition of SARS-CoV1 (HKU-39849) and SARS-CoV-2 (Ref-SKU: 026V-03883) genomes, using BWA using standard settings (*39*). Reads that mapped equally well to multiple locations were discarded. Mapping and generation of count tables was done using the MapAndGo script. Samples were normalized using RPM normalization.

Differential gene expression analysis was performed using the DESeq2 package (*40*). SARS-mapping reads were removed before analyzing the different datasets. After filtering for an adjusted p-value < 0.05, the row z-score for the 25 genes with the highest upregulation after 60 hours of exposure to SARS-CoV-2 was calculated. Data will be deposited in GEO.

The 50 genes which were most strongly induced in response to SARS-CoV-2 (padj < 0.05, ranked by fold change) were subjected to functional enrichment analysis for a biological process using ToppFun on the ToppGene Suite (https://toppgene.cchmc.org/enrichment.jsp) as described before (*41*). The 5 biological processes with highest enrichment (after FDR correction and a p-value cutoff of 0.05) for each virus are displayed with the corresponding GO term and corrected p value.

#### Transmission electron microscopy

Organoids were chemically fixed for 3 hours at room temperature with 1.5% glutaraldehyde in 0.067 M cacodylate buffered to pH 7.4 and 1 % sucrose. Samples were washed once with 0.1 M cacodylate (pH 7.4), 1 % sucrose and 3x with 0.1 M cacodylate (pH 7.4), followed by incubation in 1% osmium tetroxide and 1.5% K4Fe(CN)6 in 0.1 M sodium cacodylate (pH 7.4) for 1 hour at 4 °C. After rinsing with MQ, organoids were dehydrated at RT in a graded ethanol series (70, 90, up to 100%) and embedded in epon. Epon was polymerized for 48h at 60 °C. 60nm Ultrathin sections were cut using a diamond knife (Diatome) on a Leica UC7 ultramicrotome, and transferred onto 50 Mesh copper grids covered with a formvar and carbon film. Sections were post-stained with uranyl acetate and lead citrate.

All TEM data were collected autonomously as virtual nanoscopy slides (*26*) on FEI Tecnai T12 microscopes at 120kV using an Eagle camera. Data were stitched, uploaded, shared and annotated using Omero (*42*) and PathViewer.

Data was deposited to the Image Data Resource (https://idr.openmicroscopy.org) under accession number idr0083.”

#### Multiplex cytokine ELISA

Organoid supernatant samples, stored at −80°C, were thawed and centrifuged at 2,000 g for 5 min. The Anti-virus response panel multiplex ELISA kit (BioLegend) was used to determine cytokine concentrations in the samples using a FACSLyric (BD biosciences) flow cytometer. The procedure was performed according to the manufacturer’s instructions with the addition that capture beads were inactivated in 4% paraformaldehyde for 15 min at room temperature and washed once in wash buffer after completion of the staining protocol in a Class II Biosafety Cabinet under BSL-3 conditions.

#### Statistical analysis

Statistical analysis was performed with the GraphPad Prism 8 software. We compared differences in cytokine levels and log10 transformed viral titers by a two-way ANOVA followed by a Sidak multiple-comparison test.

### Legends Suppl figures and tables

**Supplementary figure 1.**
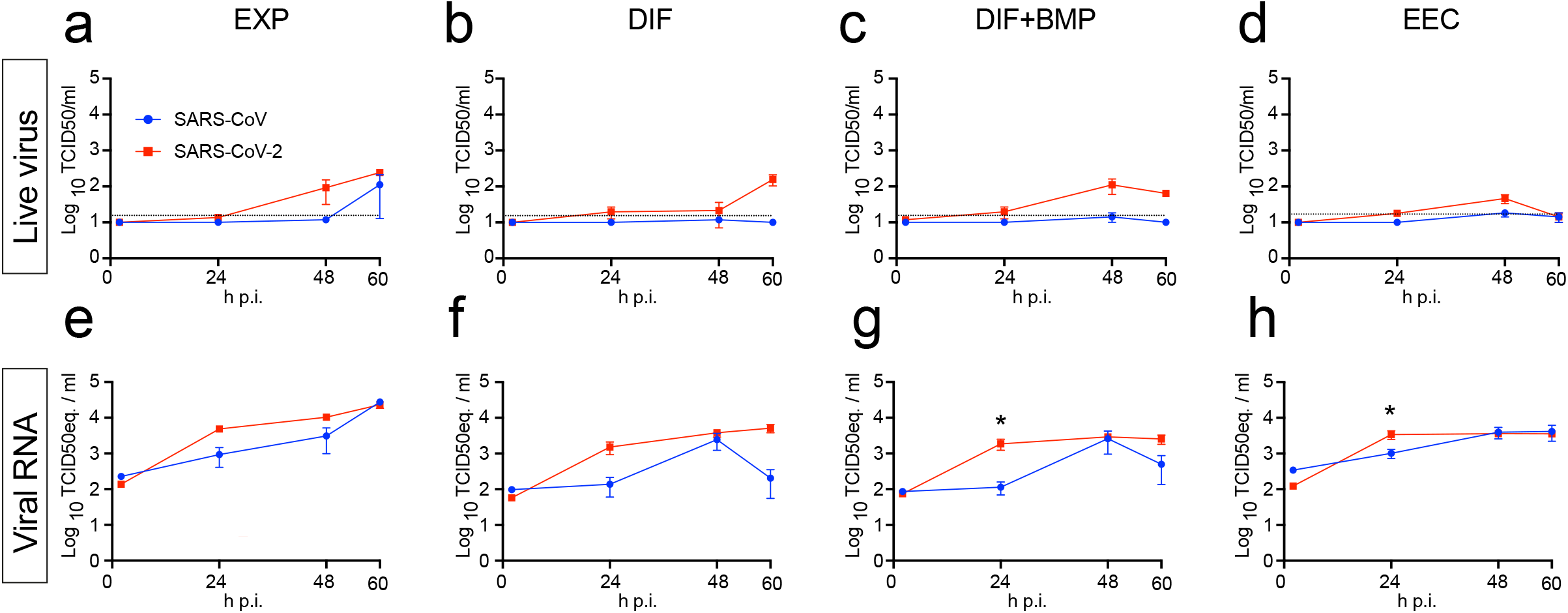
Limited secretion of SARS-CoV and SARS-CoV-2 into organoid supernatant. a-d) Live virus titers can be observed by virus titrations on VeroE6 cells of organoid supernatant at 2, 24, 48 and 60h after infection with SARS-CoV (blue) and SARS-CoV-2 (red). Different medium compositions show similar results. e-h) qPCR analysis targeting the E gene of similar timepoints and medium compositions as a-d. The dotted line indicates the lower limit of detection. Error bars represent SEM. N=3.

**Supplementary figure 2.**
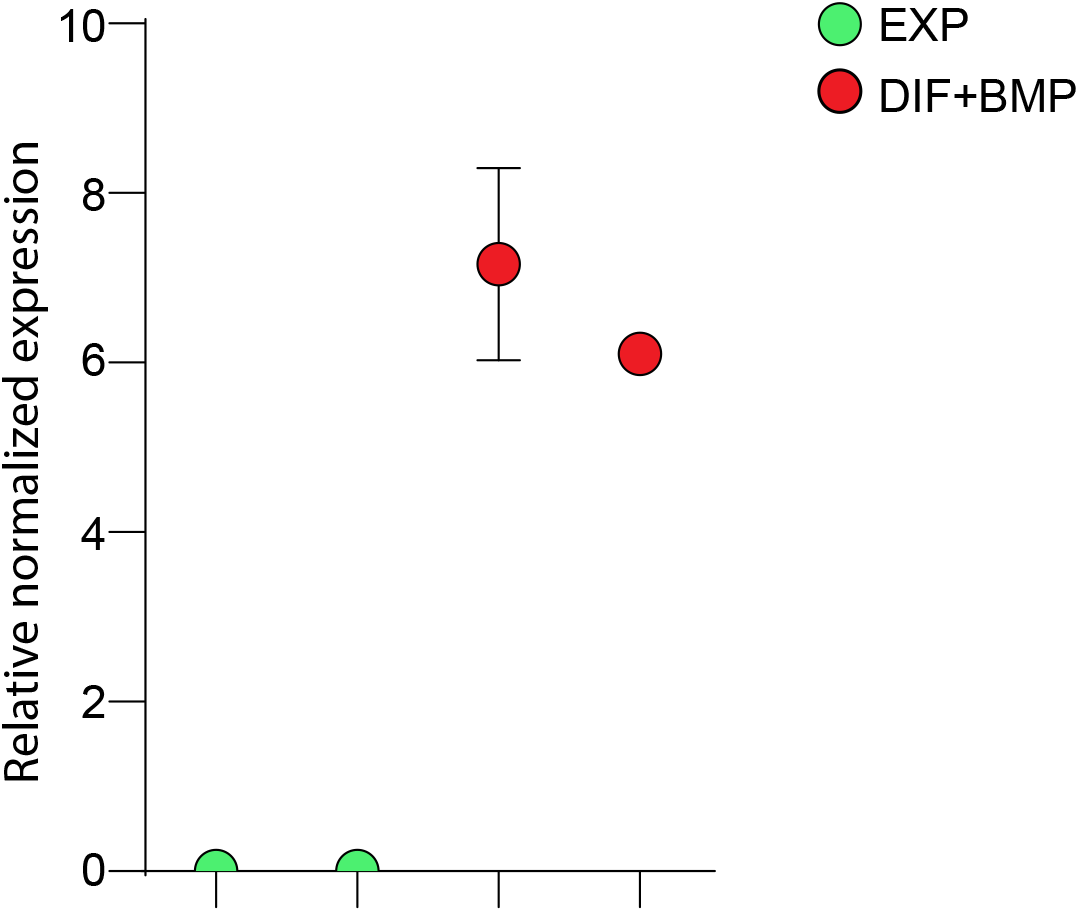
ACE2 expression is absent in proliferating organoids and increases upon BMP stimulation. qPCR analysis of ACE2 on proliferating (EXP), differentiated (DIF) without and with BMP (DIF+BMP). Expressions levels are normalized to GADPH and relative to DIF organoids.

**Supplementary figure 3.**
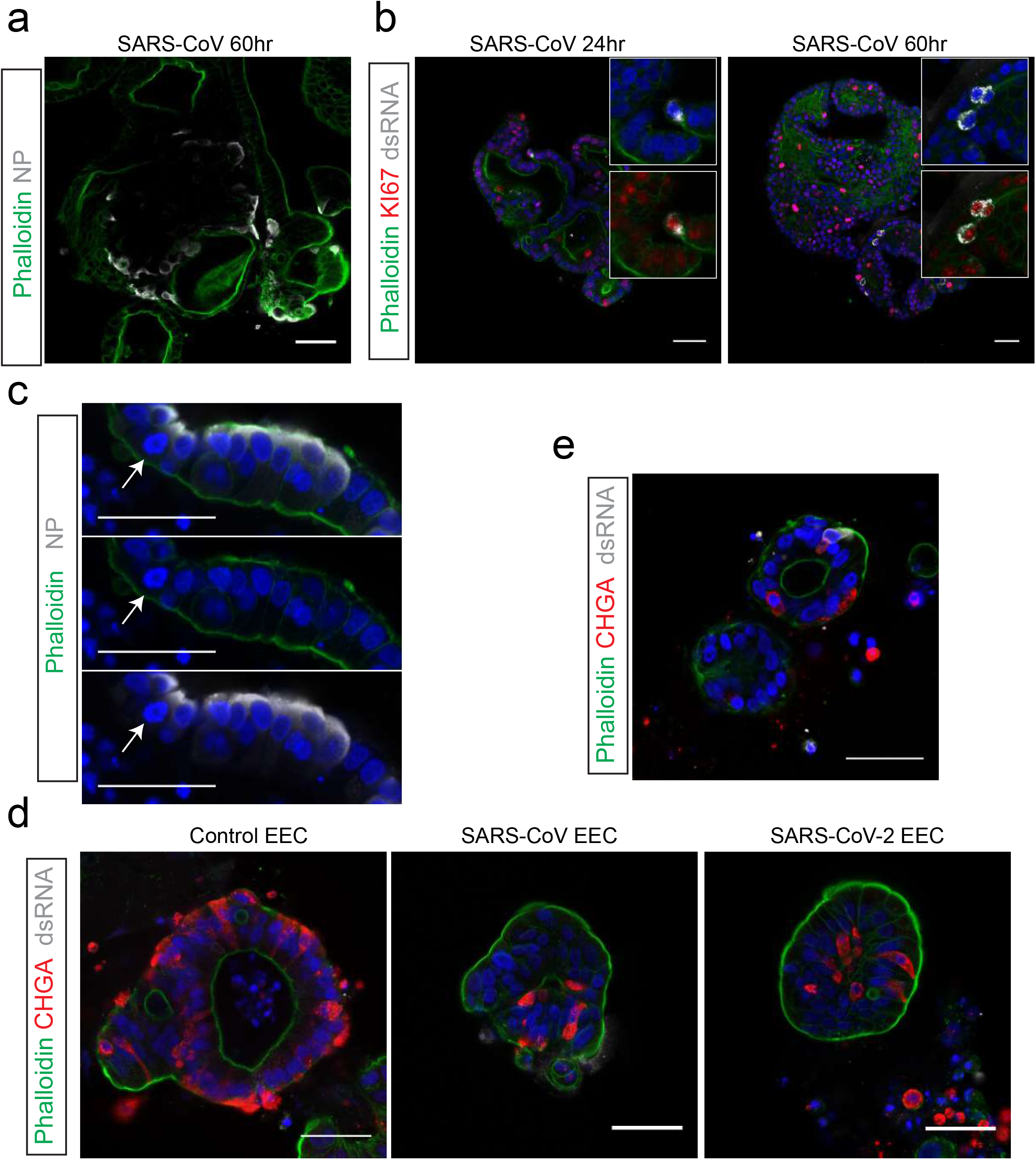
SARS-CoV infects proliferating cells but not secretory endocrine cells. a) Immunofluorescent staining of SARS-CoV-infected intestinal organoids in expansion conditions. Nucleoprotein (NP) stains viral capsid. Infected organoids contain large clusters of virus-containing cells at 60 hours after infection. b) SARS-CoV infects dividing cells that are KI67-positive. c) Example of a mitotic infected cell (arrow). d) Enteroendocrine cell (EEC) differentiated organoids do not facilitate viral infection. EECs are marked by Chromogranin A (CHGA). An average of 7 ± 5,2 CHGA+ EECs were observed per organoid section (n= 9 sections). e) Example of an EEC-differentiated organoid with a virus infected cell. All scale bars are 50 μm.

**Supplementary figure 4.**
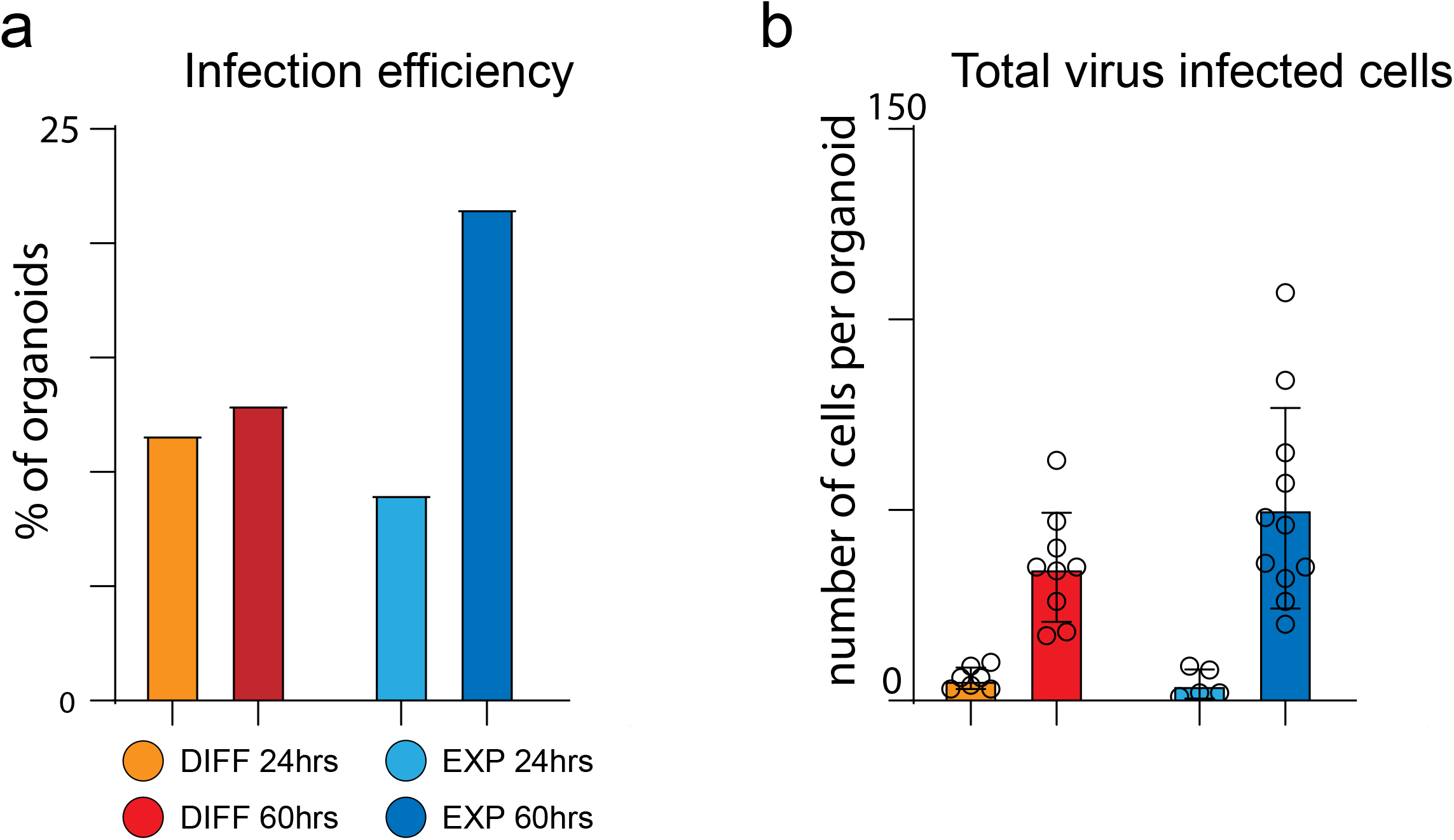
Quantification of SARS-CoV-2 infection spreading in organoids. a) Percentage of organoids harboring at least 1 SARS-CoV2 infected cell are depicted in the different culture conditions and at different timepoints. At least 50 different organoids were counted per condition. 60 hours infection in EXP medium was performed in n=2 independent experiments. b) The total number of infected cells per organoid (from a) were counted. 60 hours infection in EXP medium was performed in n=2 independent experiments.

**Supplementary figure 5.**
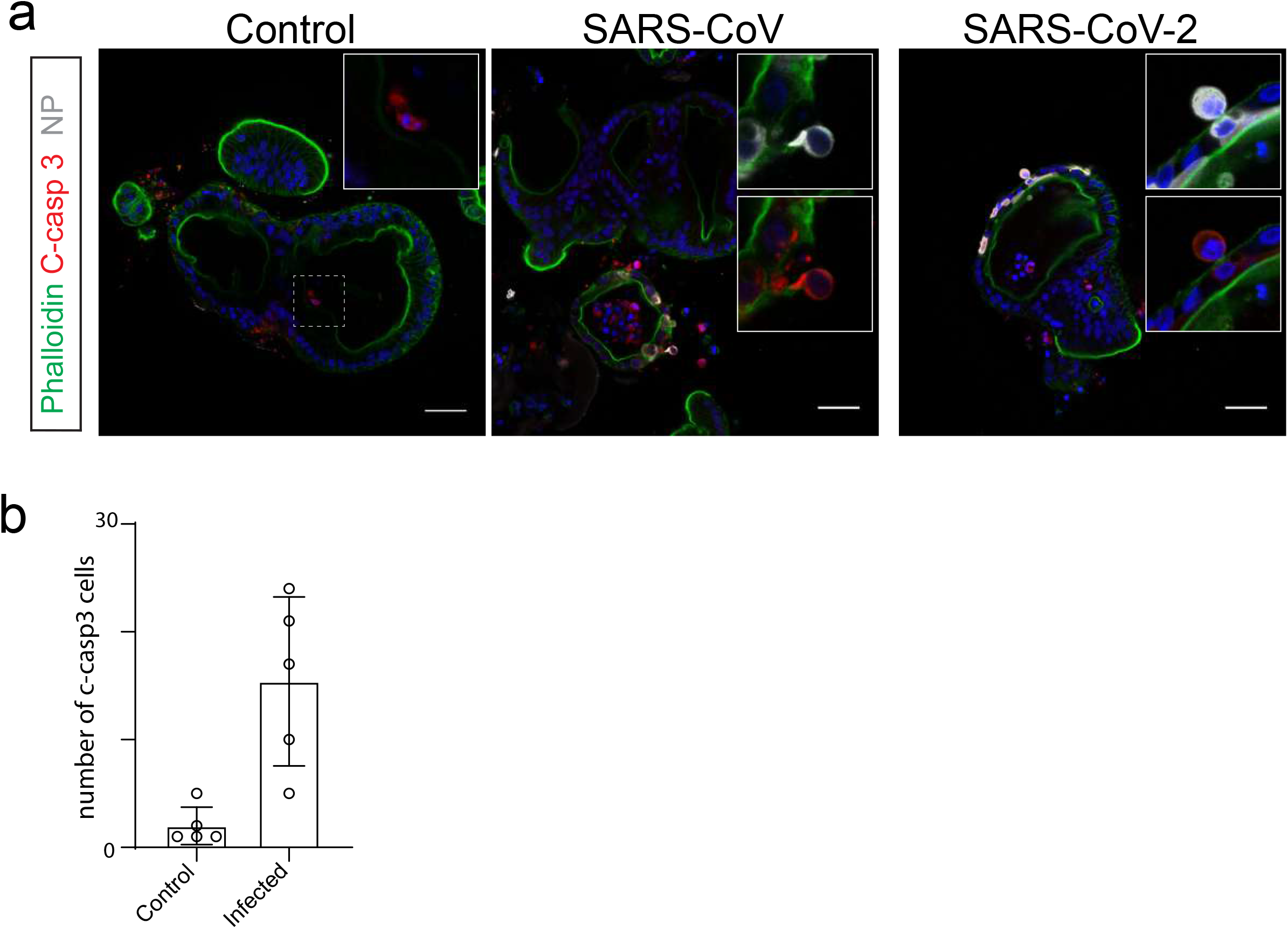
Increased apoptosis in organoids infected with SARS-CoV and SARS-CoV-2. a) SARS-CoV and −2-infected organoids show hallmarks of cell death, including detachment from the epithelial layer and cleaved-caspase 3 positivity. All scale bars are 50 μm. b) Number of cleaved-caspase 3 positive cells in control and SARS-CoV2 infected organoids were counted

**Supplementary figure 6.**
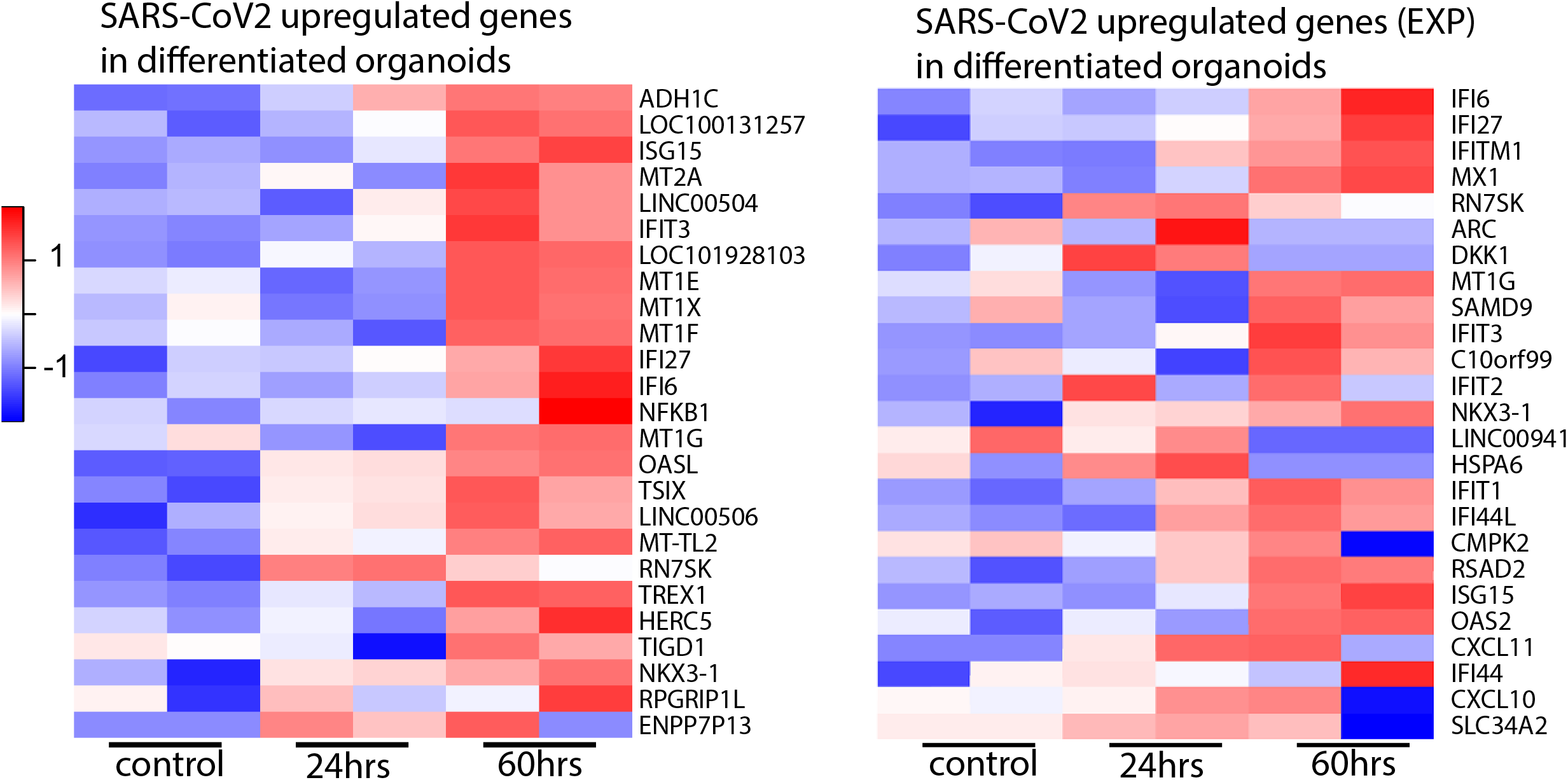
Transcriptomic analysis of SARS-CoV-2 infected differentiated organoids. Heatmaps depicting the 25 most significantly enriched genes upon SARS-CoV-2 infection in differentiated intestinal organoids (left). Right heatmap shows the top 25 signature genes from SARS-CoV2 infected EXP organoids (Figure 2a) upon SARS-CoV2 infection in differentiated intestinal organoids. Colored bar represents Z-score of log2 transformed values.

**Supplementary figure 7.**
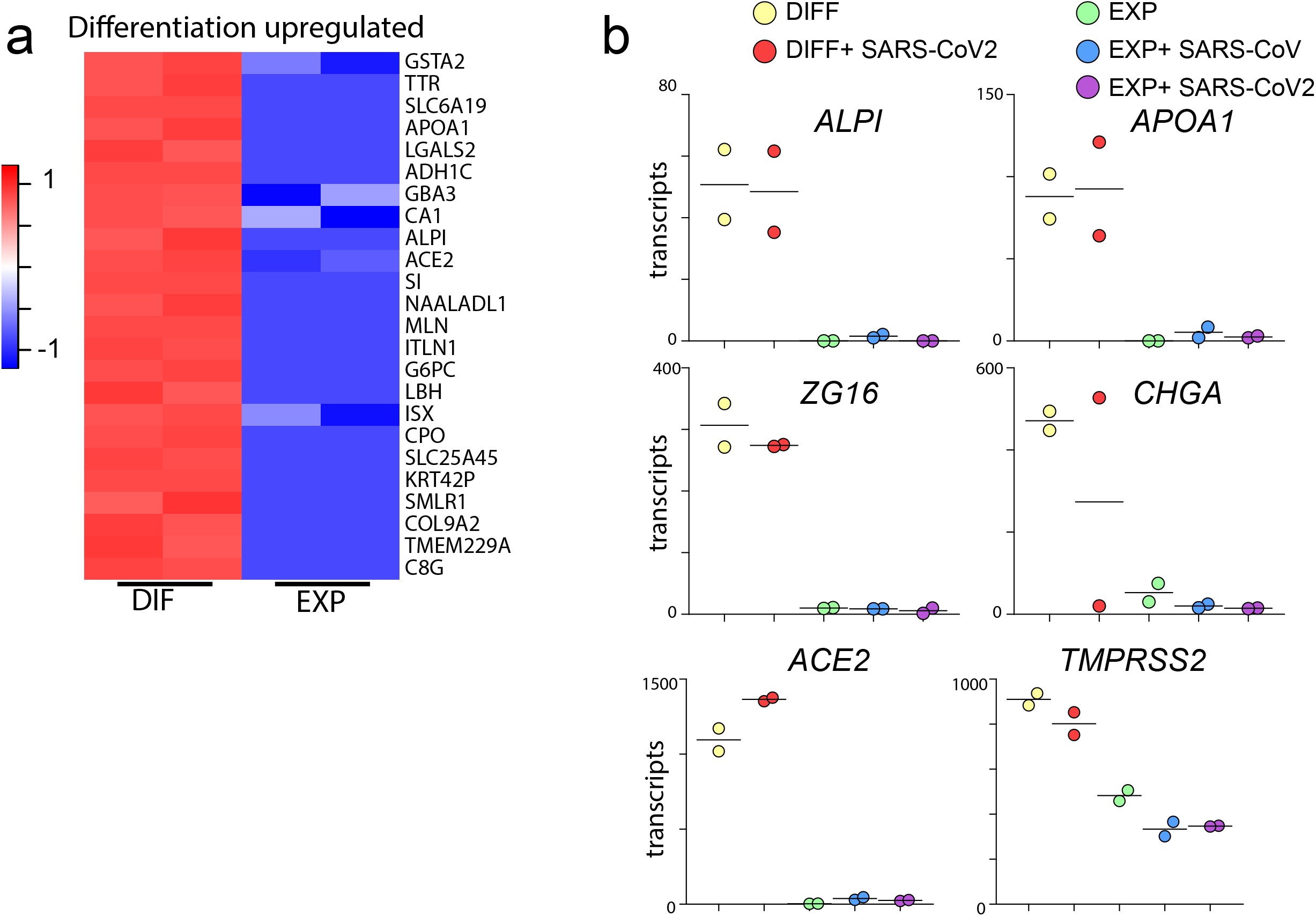
Differentiated organoids upregulate markers of mature intestinal lineages. a) Heatmaps depicting the 25 most significantly enriched genes upon differentiation compared to expansion condition. Colored bar represents Z-score of log2 transformed values. b) Graphs depicting the transcript counts of different genes enriched in mature lineages. Enterocytes (*ALPI* and *APOA1),* Goblet cells (*ZG16)* and Enteroendocrine cells (*CHGA)* increase upon differentiation. Genes required for viral entry (*ACE2* and *TMPRSS2)* are enriched in differentiated organoids.

**Supplementary figure 8.**
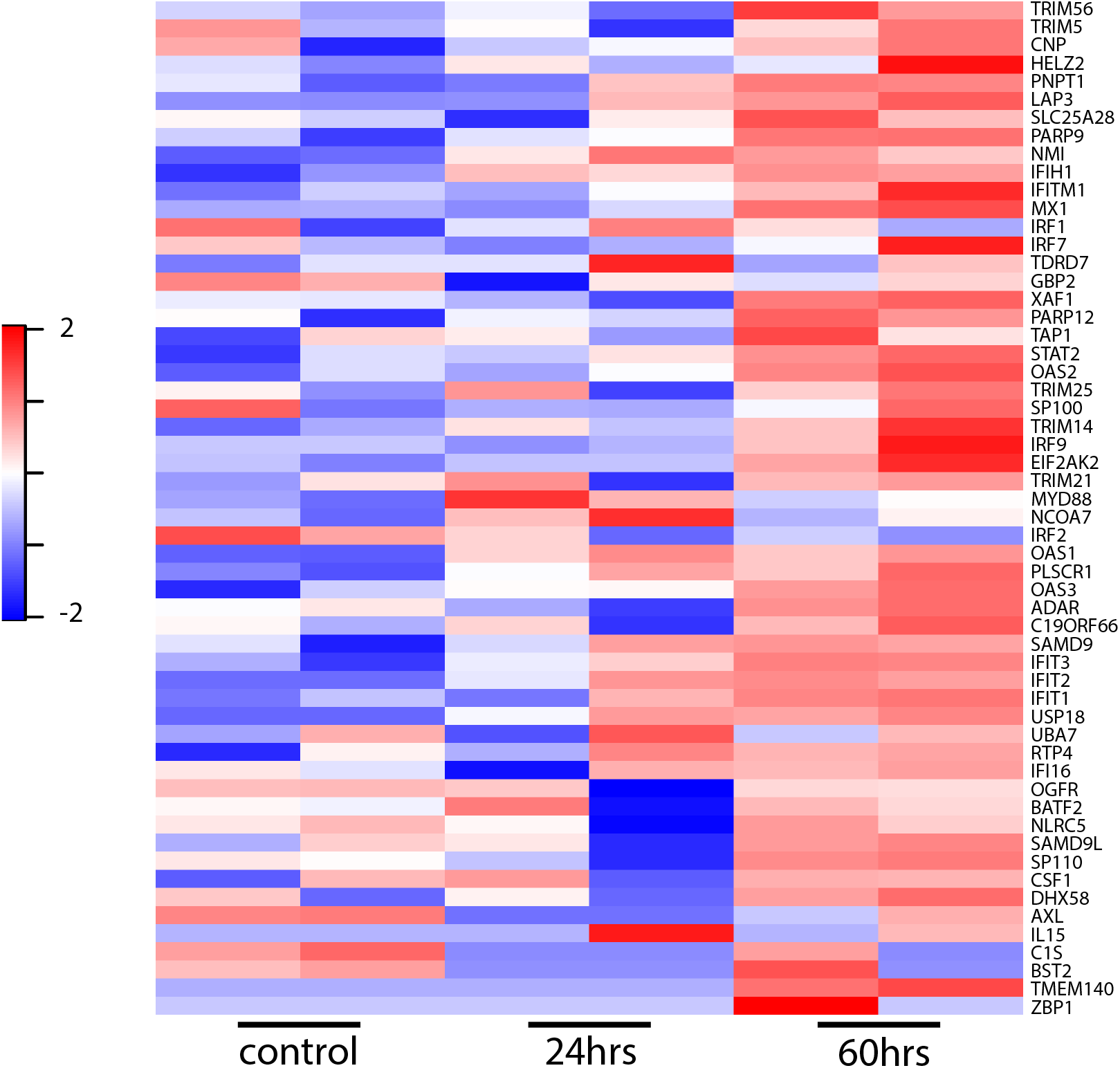
Interferon stimulated genes after SARS-CoV2 infection in intestinal organoids. Heatmaps depicting ISGs reported in (*33*) upon SARS-CoV2 infection in intestinal organoids. Colored bar represents Z-score of log2 transformed values.

**Supplementary figure 9.**
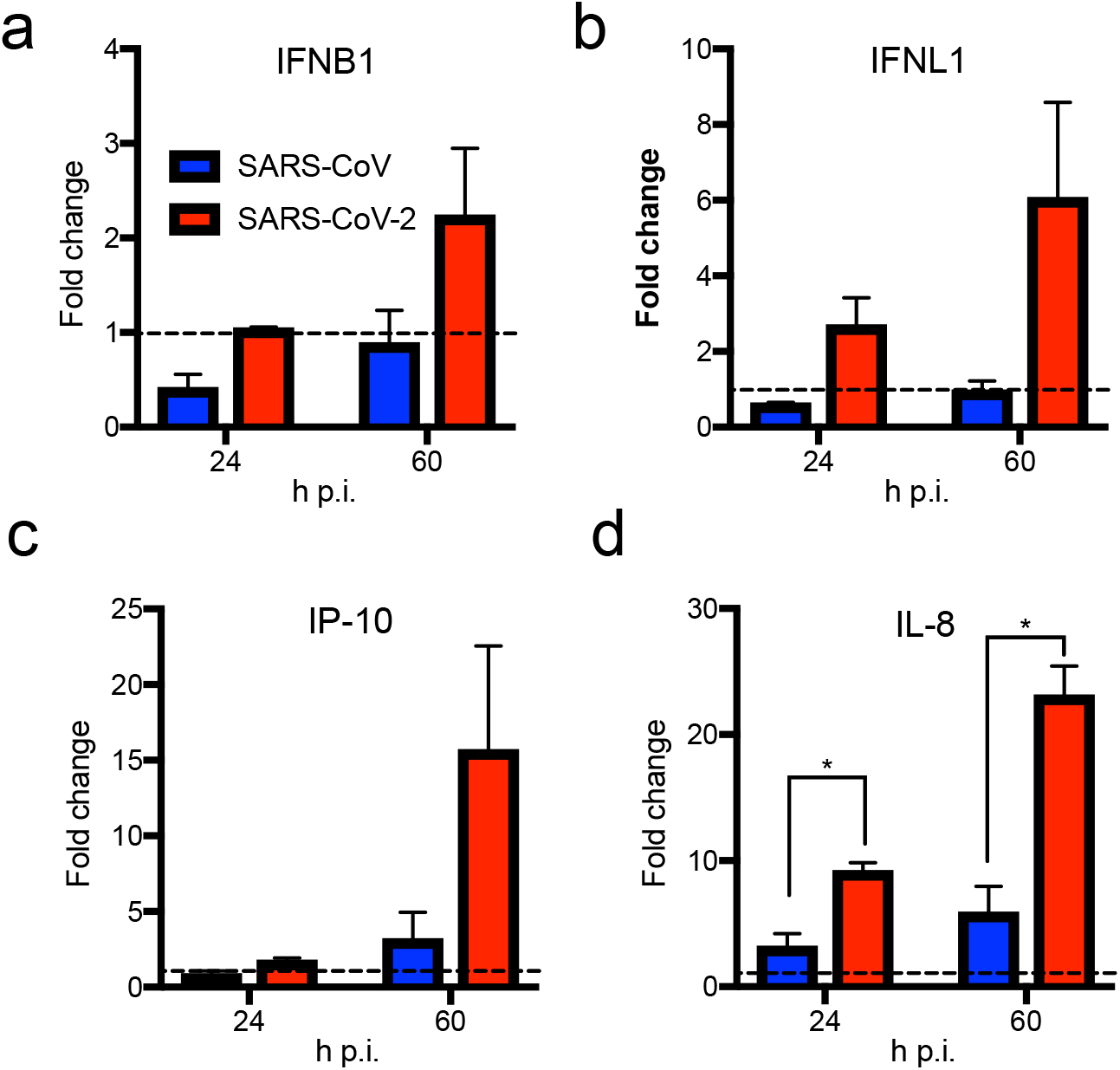
SARS-CoV-2 elicits cytokine responses in intestinal organoids. IFNB1 (a), IFNL1 (b), IP-10 (c) and IL-8 (d) levels in organoid culture supernatants were determined using Legendplex multiplex ELISA. Values were normalized to the 2h timepoint post infection. The dotted line indicates a fold change of 1. Error bars indicate SEM. N=3.

**Supplementary figure 10.**
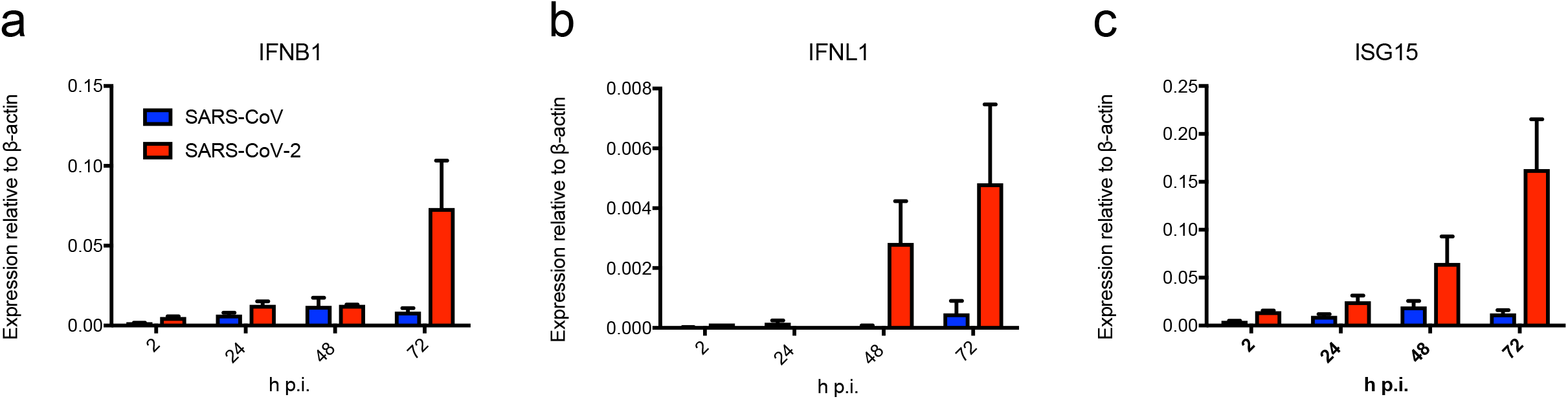
SARS-CoV-2 elicits cytokine responses in intestinal organoids as determined by qRT-PCR. IFNB1 (a), IFNL1 (b) and ISG15 (c) mRNA expression in organoids was determined by qRT-PCR. Values were normalized to β-actin expression. Error bars indicate SEM. N=3.

**Supplementary figure 11.**
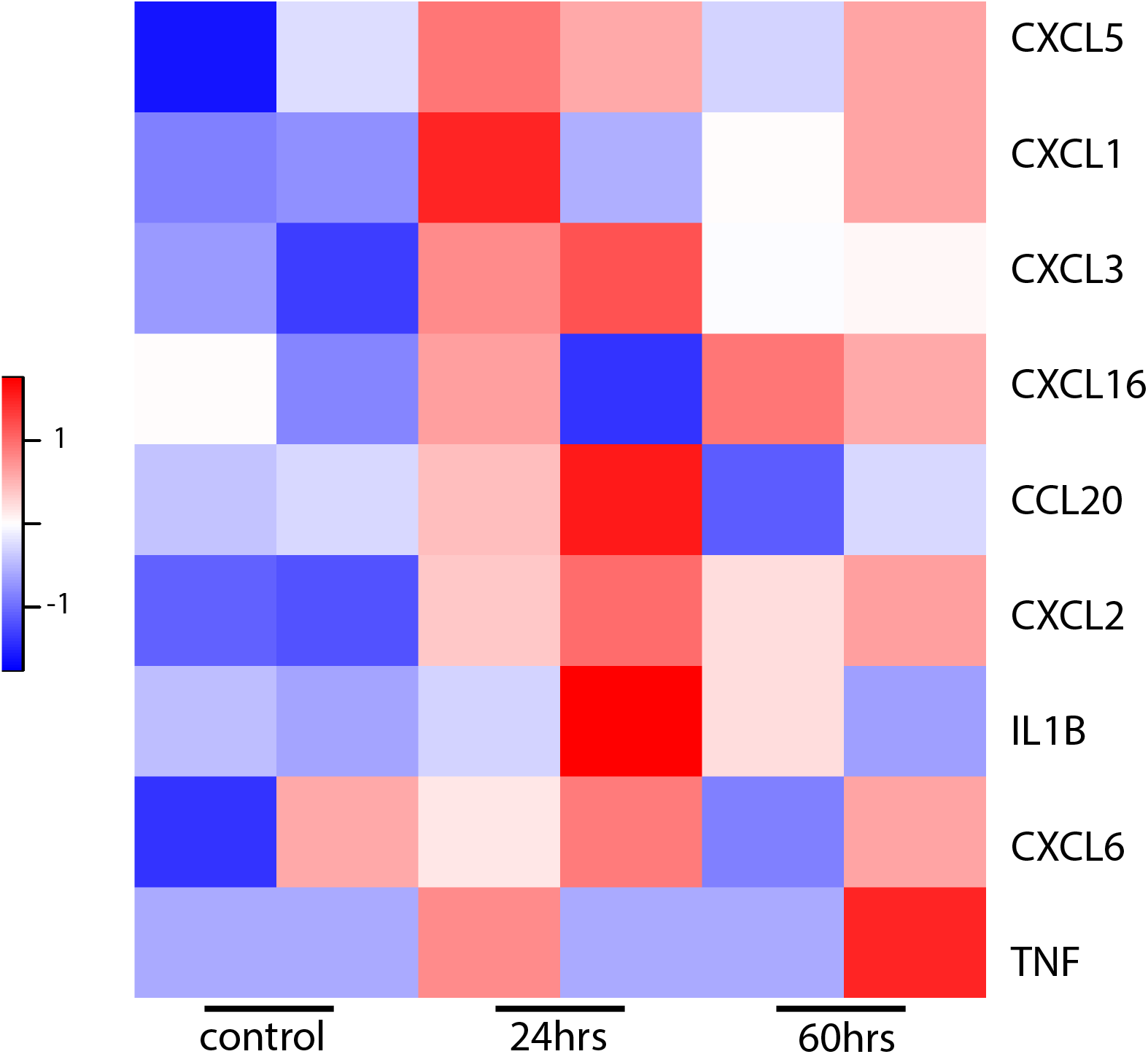
Cytokines after SARS-CoV2 infection in intestinal organoids. Heatmaps depicting virally regulated cytokines reported in (*33*) upon SARS-CoV2 infection in intestinal organoids. Colored bar represents Z-score of log2 transformed values.

**Supplementary figure 12.**
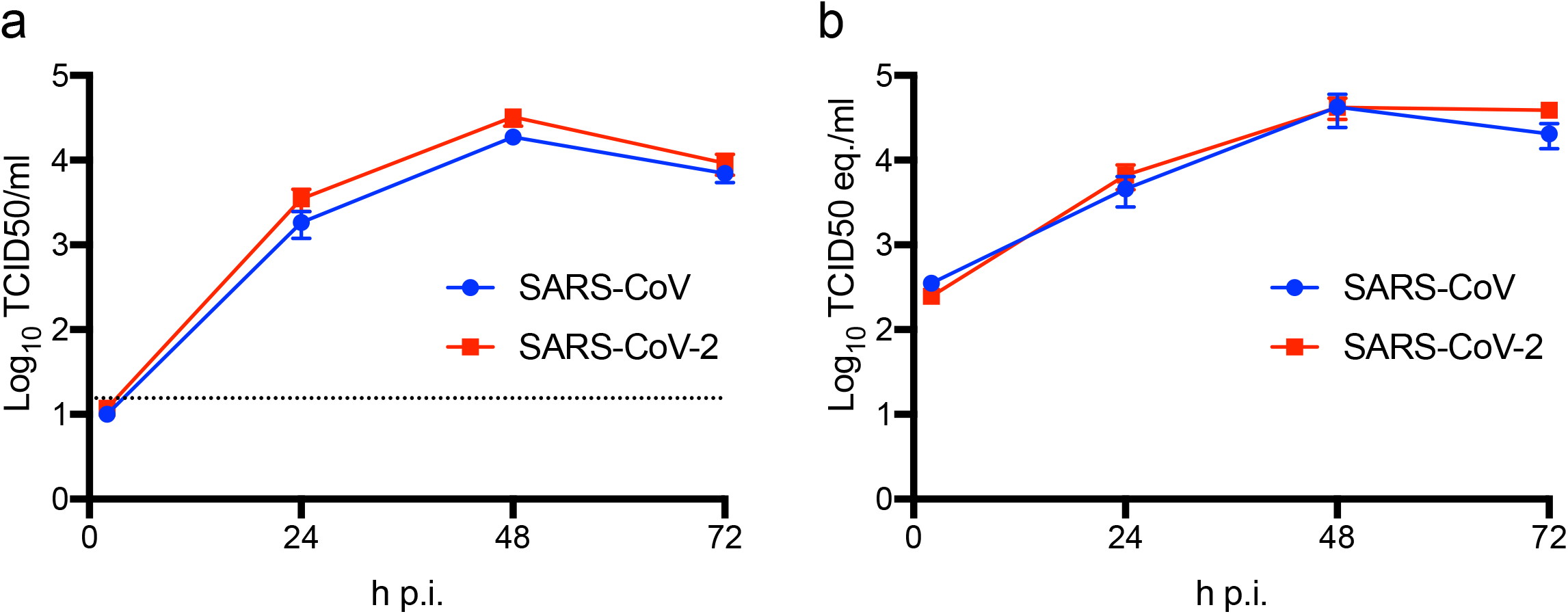
SARS-CoV and SARS-CoV-2 replicate in hSIOs in expansion medium. (a) Live virus and viral RNA (b) titers can be observed by virus titrations on VeroE6 cells of lysed organoids at 2, 24, 48 and 72h after infection with SARS-CoV (blue) and SARS-CoV-2 (red). The dotted line indicates the lower limit of detection. Error bars represent SEM. N=3.

**Supplementary figure 13.**
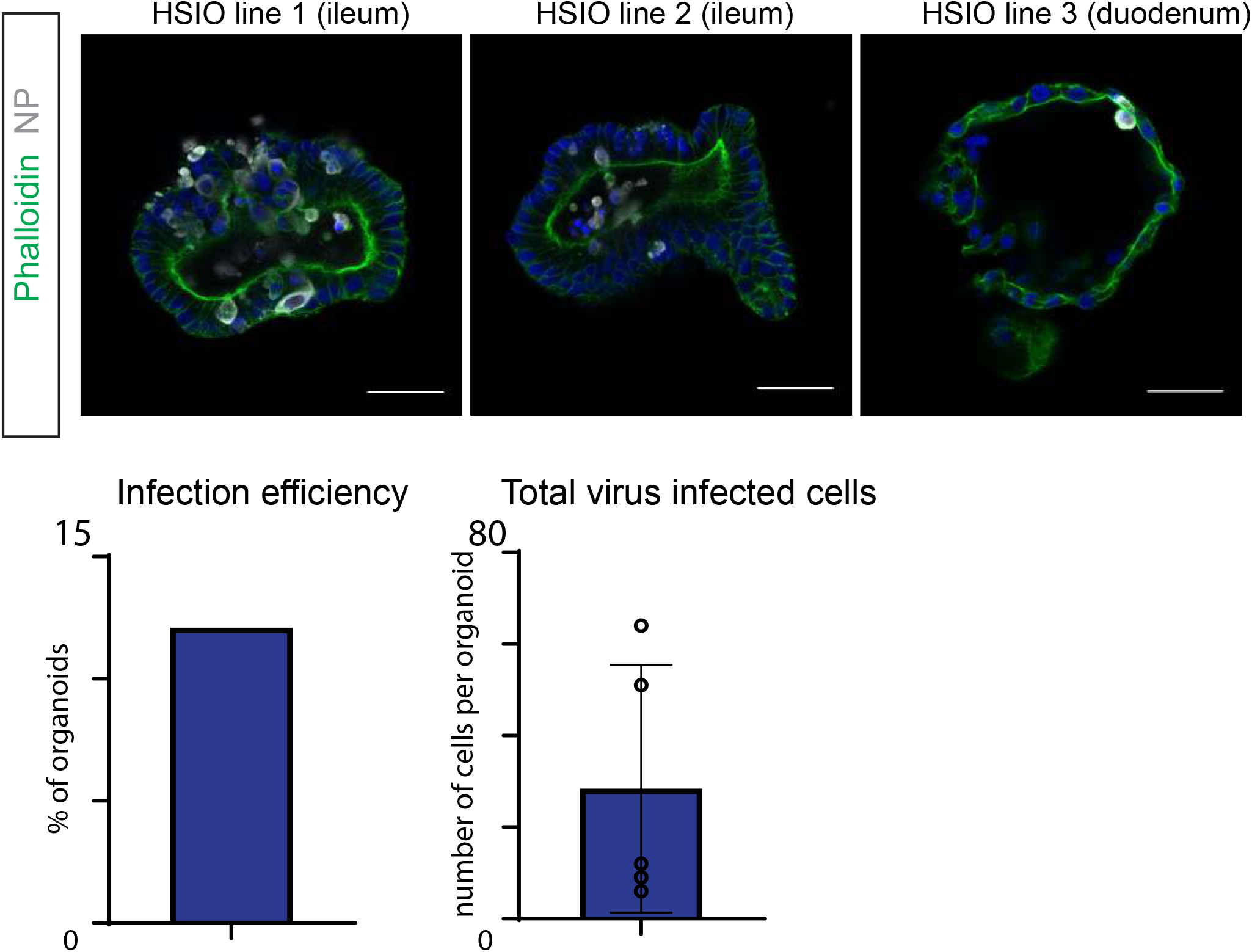
Human intestinal organoids from multiple donors are infected by SARS-CoV2. Immunofluorescent staining of SARS-CoV2-infected intestinal organoids in expansion conditions 72 hours after infection. Nucleoprotein (NP) stains viral capsid. The left panel is the same human small intestinal organoid (HSIO) line as used throughout the study, the middle panel a second line from a human ileum, and the right panel a line from a human duodenum. All scale bars are 50 μm.

**Supplementary figure 14.**
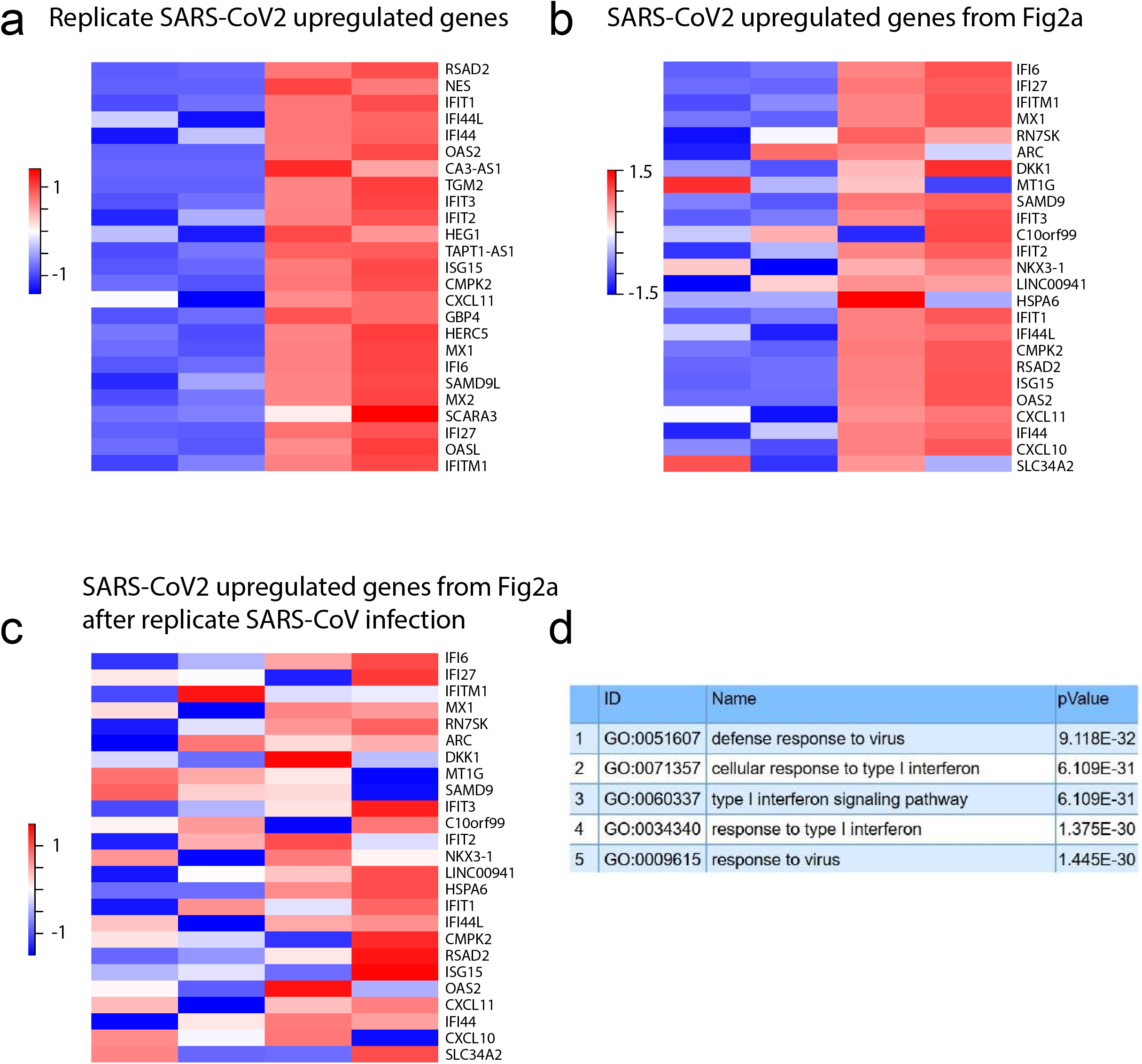
Transcriptomic analysis of a replicate SARS-CoV-2 infection in intestinal organoids. a) Heatmaps depicting the 25 most significantly enriched genes upon the replicate SARS-CoV-2 infection in expanding intestinal organoids. Colored bar represents Z - Score of log2 transformed values. b) Heatmaps depicting the 25 genes from Figure 2a in the replicate SARS-CoV-2 infected expanding organoids. Colored bar represents Z-score of log2 transformed values. c) Heatmaps depicting the 25 genes from Figure 2a in the replicate SARS-CoV infected expanding organoids. Colored bar represents Z-score of log2 transformed values. d) GO term enrichment analysis for biological processes of the 50 most significantly upregulated genes upon SARS-CoV-2 infected in intestinal organoids.

**Supplementary table 1 Normalized transcript counts in SARS-infected organoids**

**Supplementary table 2 Differentially regulated genes upon SARS-CoV2 infection in expansion organoids**

Fold change in gene expression versus control is shown after 60 hours of SARS-CoV2 infection in organoids in expansion medium.

**Supplementary table 3 Differentially regulated genes upon SARS-CoV2 infection in differentiated organoids**

Fold change in gene expression versus control is shown after 60 hours of SARS-CoV2 infection in organoids in differentiation medium.

**Supplementary table 4 Differentially regulated genes upon SARS-CoV infection in differentiated organoids**

Fold change in gene expression versus control is shown after 60 hours of SARS-CoV infection in organoids in expansion medium.

